# EmbryoTempoFormer: clip-based developmental tempo inference from zebrafish brightfield time-lapse microscopy

**DOI:** 10.64898/2026.03.09.710433

**Authors:** Li-Jia-Yu Deng, Pei-Ran Lin, Luo-Tong Xie

## Abstract

Nominal hours post fertilization (hpf) are widely used to index zebrafish embryogenesis, yet under condition shifts—such as temperature change, genetic perturbation, or environmental stress—nominal time can decouple from true developmental progression. In such settings, biologically meaningful variation is better described as a systematic change in developmental tempo rather than a simple temporal offset. Here we introduce an embryo-resolved framework that treats developmental tempo as the primary quantity of interest in brightfield time-lapse imaging. We present EmbryoTempoFormer (ETF), a clip-based CNN–Transformer that predicts developmental progression from short time-lapse clips and is trained with a within-embryo temporal-difference consistency regularizer to promote temporally coherent trajectories. Crucially, we couple model predictions with an embryo-level inference and statistical workflow: temporally correlated clip-level outputs are aggregated into interpretable embryo-level tempo and stability readouts, and cross-condition effects are quantified using embryo-bootstrap confidence intervals with embryos—rather than frames or clips—as independent units, avoiding pseudo-replication. Using temperature perturbation as a representative domain shift, we robustly quantify condition-induced changes in global developmental dynamics and show that developmental delay predominantly manifests as reduced developmental tempo. This framework enables statistically principled, high-throughput phenotyping for perturbation screens, drug assays, and environmental stress studies.

**HIGHLIGHTS:**

- Clip-based CNN–Transformer predicts developmental time from brightfield time-lapse microscopy.
- Within-embryo temporal-difference consistency improves trajectory self-consistency.
- Embryo-level anchored tempo slopes enable interpretable cross-condition comparisons.
- Reproducible pipeline via code, scripts, and a Zenodo bundle with embryo-level inference

**Graphical abstract:** 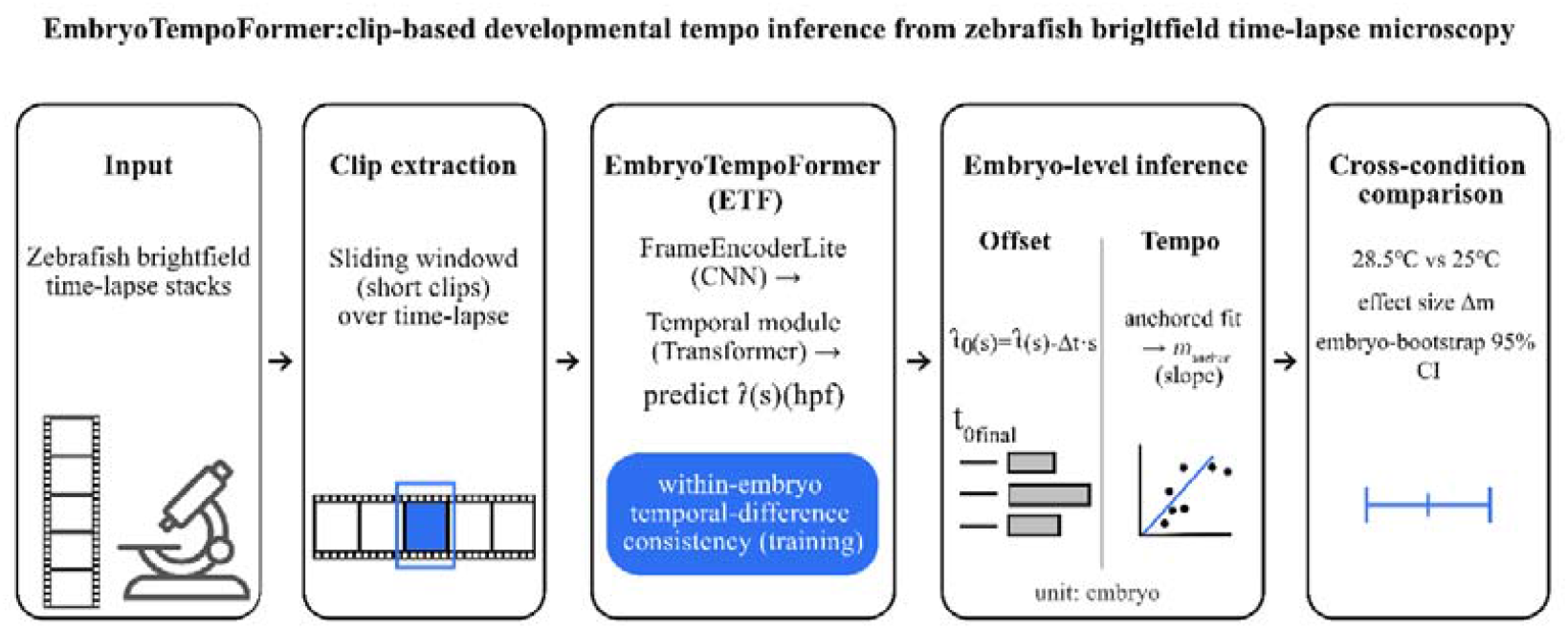

## 1. Introduction

Zebrafish (*Danio rerio*) are widely used in developmental biology, genetics, toxicology, and drug screening due to their transparent embryos, rapid development, and compatibility with high-throughput perturbation experiments. Accurate staging is fundamental for aligning phenotypes across time and experimental conditions. Traditionally, staging combines nominal hours post fertilization (hpf) with morphological criteria summarized in standard staging guides [1]. In practice, however, investigators are often concerned not only with whether a specific stage has been reached, but also with whether genetic perturbations, drug treatments, or environmental changes induce developmental delay or acceleration and how large such effects are at the population level. Manual staging is therefore both labor-intensive and subject to inter-observer variability, limiting stable quantification on large-scale imaging data.

A further challenge is that nominal hpf is not a universal measure of developmental progress. Even under standard temperature (28.5°C), hpf is only an approximation; under condition shifts—most notably temperature—developmental dynamics can change systematically, and developmental progression can vary across embryos and conditions, making nominal hpf an imperfect proxy for developmental state [2, 3]. In such settings, developmental delay is not necessarily a purely additive shift in time; rather, developmental tempo (speed and rhythm) itself may change, potentially in a stage-dependent and non-linear manner. Consequently, using nominal time alone often fails to provide an interpretable and statistically valid quantification of tempo changes induced by altered conditions.

Recent machine learning approaches have made automated zebrafish analysis scalable. Earlier work used handcrafted features and conventional classifiers to assign embryos to discrete stages, demonstrating feasibility while remaining sensitive to feature engineering and imaging conditions [4]. More recently, deep learning methods such as KimmelNet modeled staging as continuous regression by predicting developmental time directly from single 2D brightfield images, enabling automated staging and population-level delay detection [5]. Parallel work emphasized developmental time and tempo as objects of inference, learning relationships among developmental states to reconstruct time mappings across conditions and revealing non-linear tempo characteristics across developmental phases [6]. These advances suggest that under systematic condition shifts (e.g., temperature changes), nominal hpf may be an unstable reference and an inappropriate external “accuracy” target. At the same time, the increasing availability of brightfield time-lapse microscopy introduces a practical inferential pitfall that can undermine statistical conclusions if overlooked. In practice, long sequences are often processed via densely sampled frames or overlapping sliding windows (clips) for training or inference. Predictions obtained from the same embryo are strongly correlated; treating such correlated windows as independent samples for evaluation, hypothesis testing, or uncertainty estimation constitutes pseudo-replication and can lead to overconfident conclusions [7]. This issue becomes particularly pronounced under condition shifts, where nominal time is no longer a stable reference and window-level sample inflation can further distort downstream comparisons. Therefore, a robust framework for time-lapse analysis should (i) exploit temporal information, (ii) promote coherent within-embryo trajectories, and (iii) perform inferential statistics at the embryo level.

Here we propose the EmbryoTempoFormer (ETF) model, a clip-based approach for zebrafish brightfield time-lapse microscopy that predicts developmental time from short clips and enforces within-embryo temporal coherence during training via a temporal-difference consistency regularizer. In a downstream inference and statistical workflow, correlated clip predictions within each embryo are aggregated into embryo-level tempo readouts suitable for cross-condition comparison, with embryos treated as the independent statistical units to avoid pseudo-replication [7]. This embryo-resolved framing is compatible with the broader trend toward time-resolved phenotyping tasks such as developmental event detection, which aim to localize temporal phenotypes from microscopy sequences [8]. The main contributions of this work are:

1. A clip-based CNN–Transformer framework for predicting developmental time from brightfield time-lapse microscopy.
2. A within-embryo temporal-difference consistency regularizer that improves trajectory self-consistency across overlapping predictions.
3. Embryo-level tempo readouts based on an anchored slope (m_anchor) and residual-based stability metrics for interpretable cross-condition comparisons.
4. A reproducible end-to-end pipeline and statistically rigorous cross-condition inference using embryo-bootstrap confidence intervals with embryos as the independent statistical units.

## 2. Related work

### 2.1. Automated zebrafish staging from images

Early automated staging approaches used handcrafted features and conventional machine learning models to assign embryos to discrete stages, demonstrating feasibility while remaining sensitive to feature engineering and imaging variability [4]. More recently, deep learning has enabled continuous staging by regressing developmental time from single 2D brightfield images. KimmelNet is a representative example that achieves high agreement with expert staging and supports population-level detection of overall delay or acceleration under standard conditions [5]. While effective, single-frame regression does not explicitly exploit temporal context from time-lapse microscopy and does not by itself resolve inferential issues that arise when dense sequences are analyzed using correlated frames or overlapping windows.

### 2.2. Developmental time and tempo under condition shifts

Beyond absolute time regression, recent work emphasized developmental time and tempo as objects of inference. By learning developmental state similarity and reconstructing time mappings across conditions, such approaches capture non-linear characteristics of developmental progression and reveal tempo changes across developmental phases [6]. This perspective is particularly relevant under systematic condition shifts such as temperature changes, where nominal hpf may be an unstable alignment reference and “accuracy against a nominal clock” can be conceptually inappropriate. Our work is complementary: we retain clip-to-time regression as a practical primitive while focusing on embryo-level tempo readouts and statistically valid comparisons under correlated time-lapse sampling.

### 2.3. Deep learning for zebrafish phenotyping beyond staging

Deep learning has been applied to a broad range of zebrafish imaging tasks beyond staging, including attention-based phenotype recognition for high-throughput screening, deformation classification, robustness-oriented multi-stage pipelines for brightfield larvae, anomaly detection of abnormal development, integration with microfluidics for real-time and non-invasive embryo handling, and linking embryo phenotypes to mechanistic signaling pathways [9-14]. Reviews summarize this rapid expansion and highlight persistent challenges in interpretability, robustness, and generalization. Time-resolved phenotyping tasks have also expanded toward developmental event detection, where models identify the timing of specific developmental events from imaging sequences [8, 15]. These directions underscore the growing availability of dense imaging data and the need for principled temporal readouts that remain interpretable under condition shifts.

### 2.4. Time-lapse evaluation and pseudo-replication

Time-lapse sequences are frequently processed through densely sampled frames or overlapping sliding windows to enable learning and inference over long recordings. However, windows from the same embryo are strongly correlated; treating them as independent samples inflates effective sample size and constitutes pseudo-replication in statistical inference [7]. This pitfall is model-agnostic and becomes particularly consequential when downstream analyses report uncertainty or significance using window-level samples rather than embryo-level units. ETF addresses this issue by enforcing within-embryo temporal coherence during training and by performing inferential statistics at the embryo level.

## 3. Methods

### 3.1. Processing of OME-format microscopy data and NumPy preprocessing pipeline

Raw zebrafish brightfield time-lapse stacks were loaded as multi-page sequences using tifffile and represented as a NumPy array of shape [T, H, W]. For end-to-end reproducibility, we distribute the processed arrays in the Zenodo bundle. Each stack was converted into a fixed-shape processed array uint8 [T, H, W] by:

1. Percentile clipping and normalization. Intensities were clipped at percentile bounds (*p*_*lo*_, (*p*_*hi*_) (default: 1–99), linearly mapped to [0, 1], then scaled to [0, 255] and stored as 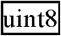.
2. Resizing. Frames were resized to 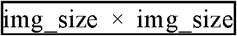 (default: 384 × 384) using bilinear interpolation (PIL).
3. Time padding/trimming. The temporal axis was standardized to 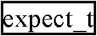 frames (default: *T* = 192).
4. Storage. One processed 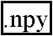 file per embryo; embryo IDs correspond to file stems.

Processed arrays were accessed via NumPy memmap 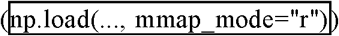 to enable efficient random access during training and evaluation.

### 3.2 Time base, clip definition, and sliding-window inference

We use the dataset timing constants *T*0 = 4.5 hpf and Δ*t* = 0.25 h/frame (15 min). A clip is defined as a contiguous window of length *L* frames extracted from a full embryo sequence:

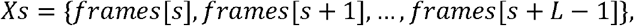

*s* is the clip start index and *L*= 24 by default. The nominal mapping from start index to nominal time is:

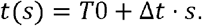

During inference, we apply sliding windows with stride *k*= 8, yielding correlated clip predictions within each embryo. Embryos (not clips/windows) are treated as the independent statistical units for inferential analyses to avoid pseudo-replication [7].

### 3.3. Dataset splits, exclusions, and evaluation sets

All dataset splits are embryo-level (one embryo corresponds to one processed time-lapse stack). We evaluate on two embryo-level test sets:

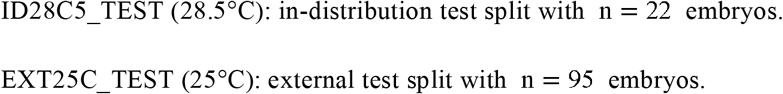

The 25°C split file lists 96 processed embryos, with 1 embryo excluded by a fixed filename rule (excluded: 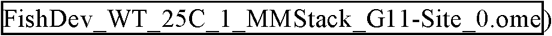, leaving 95 test embryos. For the 28.5°C condition, the dataset comprised 192 raw embryo stacks (two 96-well plates); 51 stacks were excluded following predefined filtering rules (nonviable embryos and/or embryos outside the field of view), leaving 141 included embryos that were split into train/val/test as 98/21/22. Exclusion criteria follow the original KimmelNet preprocessing pipeline and were additionally verified by manual review; no exclusions were performed to selectively improve model performance [4]. These counts are summarized in Table 1.

**Table 1.** Embryo-level dataset splits and exclusions.

| Condition<br>(temperature) | Split type | Raw<br>embryos<br>(pre-filter) | Excluded<br>embryos | Included<br>embryos | Train<br>embryos | Val<br>embryos | Test<br>embryos |
| --- | --- | --- | --- | --- | --- | --- | --- |
| 25°C<br>(EXT25C) | test-only<br>split | 96 | 1 | 95 | 0 | 0 | 95 |
| 28.5°C<br>(ID28C5) | train/val/test<br>split | 192 | 51 | 141 | 98 | 21 | 22 |

### 3.4. Training and evaluation datasets

Training uses within-embryo paired sampling: each training sample consists of two clips 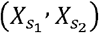 extracted from the same embryo sequence. This pairing supports the within-embryo temporal-difference consistency objective (Section 3.7). To improve sampling diversity, clip start indices are jittered within a small range (default: 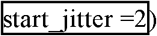. Evaluation uses a deterministic sliding-window protocol that enumerates all valid clip start indices for each embryo at a fixed stride (Section 3.2). Because sliding windows from the same embryo are strongly correlated, window-level metrics computed over all clips are reported for descriptive comparison only; inferential analyses treat embryos as the independent statistical units to avoid pseudo-replication [7].

### 3.5. Data augmentation (training only)

During training, we apply optional clip augmentations defined by 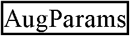. Spatial transforms (e.g., horizontal flip and affine transforms) are applied consistently across frames within a clip to preserve temporal coherence. We additionally use intensity perturbations (gamma/contrast/brightness), shading, additive noise, blur, and limited random frame drop. Augmented clips are clipped to [0, 1] before being passed to the model.

### 3.6. Model architecture (EmbryoTempoFormer, ETF; full model)

EmbryoTempoFormer (ETF) predicts a scalar developmental time 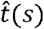 (hpf) from a grayscale clip *X*_*s*_. For a batch of clips, the input tensor has shape [*B, L*, 1, *H, W*] (default *L* = 24).

Frame encoder (FrameEncoderLite): Each frame is encoded independently into a token using a lightweight depthwise-separable CNN with GroupNorm and squeeze-and-excitation blocks [16-18]. The encoder outputs one token per frame, followed by global average pooling and a linear projection to token dimension *D* = 128. For efficiency, frame encoding is performed in chunks 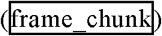 and optionally uses gradient checkpointing for memory control. For memory control, frame encoding is performed in temporal chunks 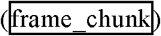 and can use chunk-level gradient checkpointing of the frame encoder 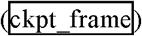, controlled by the selected memory (profile 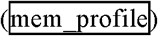.

Temporal aggregation: Temporal aggregation is controlled by 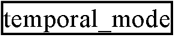, yielding four model variants used in this study:

1. cnn_single: single-frame baseline; a single frame token is used for regression (no temporal context).
2. meanpool: uniform temporal aggregation by averaging frame tokens (followed by normalization).
3. nocons: temporal Transformer aggregation without the temporal-difference consistency term (i.e., ii. *λ*_diff_ =0 in Section 3.7).
4. full (ETF): temporal Transformer aggregation with the temporal-difference consistency term enabled (ii. *λ*_diff_ = 1).

For Transformer-based variants (nocons and full/ETF), a learnable CLS token is prepended to the frame-token sequence and processed by a stack of Transformer encoder blocks with rotary positional embeddings (RoPE) applied to queries/keys [19, 20]. The final CLS token is used for regression. Training optionally applies temporal token dropout across frame tokens with rescaling to preserve expected magnitude.

Regression head: The regression head is Linear(*D* → *D*) → SiLU → Dropout _→_ Linear(D → 1).

The number of trainable parameters is 0.276M for cnn_single/meanpool and 0.804M for nocons/full.

### 3.7. Training objective and optimization

Training samples are constructed by within-embryo paired sampling: two clips 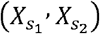 are drawn from the same embryo, with clip start indices *s*_1_ and *s*_2_. The model outputs clip-level time predictions 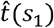 and 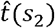 The nominal target time for a start index *s* is *t*(*s*) = *T*0 + Δ*t* · *s* (Section 3.2).

Absolute regression loss. We supervise each clip prediction against its nominal target using an elementwise regression loss *ℓ* (·, ·) (L1 or SmoothL1/Huber) [21]:

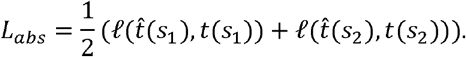

Within-embryo temporal-difference consistency. To encourage coherent within-embryo temporal progression across overlapping windows, we add a temporal-difference consistency term that constrains the predicted time difference between two clips from the same embryo to match the known sampling interval:

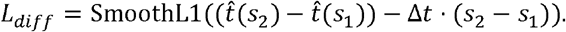

Total loss with ramp-up: The overall training objective is:

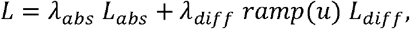

where *ramp*(*u*) ∈ [0,1] increases linearly from 0 to 1 over the first 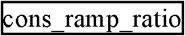 fraction of optimization steps (default 0.2). In the nocons ablation, *λ*._*diff*_ = 0; in the full (ETF) model, *λ*._*diff*_ = 1.

Optimization: We train all variants with AdamW and a cosine learning-rate schedule; all hyperparameters are reported in Section 3.8 [22].

### 3.8. Training configuration (EXP4; stored in checkpoint cfg)

Released checkpoints store the full training configuration in 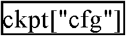 (also included in the Zenodo reproducibility bundle). Unless otherwise noted, all model variants were trained under the same configuration; the only ablation-dependent setting is 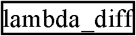 (0 for nocons, 1 for full (ETF); Section 3.7).

Data and sampling. We used 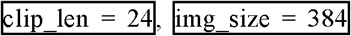, and 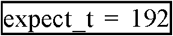. Models were trained for 300 epochs with 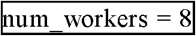. Training uses within-embryo paired sampling, drawing 32 paired clip pairs per embryo per epoch with 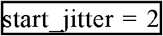. Dataset caching used 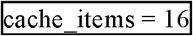.

Batching (DDP). We used 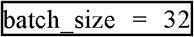 as the per-GPU batch size under distributed data parallel (DDP), and 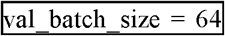 batch per GPU. (The effective global batch size scales with 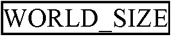 and 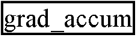, both stored in 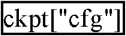.)

Optimization and scheduling. Optimization used AdamW with 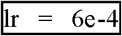 and 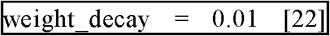. The learning-rate schedule applied linear warm-up 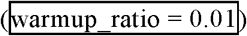 followed by cosine decay to 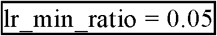.

Model hyperparameters. For Transformer-based variants, we used 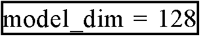, 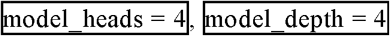, and 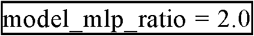. The CNN frame encoder used 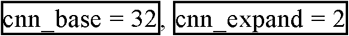, and 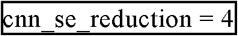.

Loss settings and training stabilization. We used 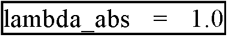 with 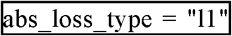 and a consistency ramp-up of 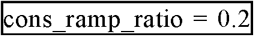 (Section 3.7). Training used mixed precision 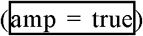 and exponential moving average evaluation with 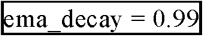 and 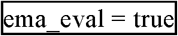.

Reproducibility and memory profile. The random seed was 42. To mitigate GPU out-of-memory errors at the reported resolution and batch size, we used a memory-optimized configuration 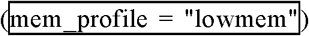, corresponding to chunked frame encoding 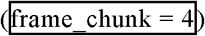 with chunk-level frame-encoder checkpointing enabled 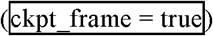 as implemented in the released code.

### 3.9. Embryo-level inference and tempo estimation

Sliding-window inference yields clip-level time predictions 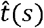 (hpf) at multiple window start indices *s* within each embryo (Section 3.2). Because overlapping clips within an embryo are strongly correlated, we summarize predictions into embryo-level readouts.

Optional start-time offset diagnostic (descriptive only): Sliding-window inference yields clip-level time predictions 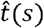 at multiple window start indices *s* within each embryo (Section 3.2). Because fertilization/collection is not perfectly synchronized, the effective time zero can vary across embryos. To provide a descriptive diagnostic of this start-time uncertainty, for each start *s* we compute an intercept-like quantity

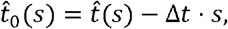

where Δ*t* is the sampling interval. We optionally summarize 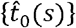 within an embryo using a trimmed mean to obtain *t*0_final_ (trim proportion = 0.2, i.e., discarding the lowest and highest 20% of values). This offset summary is reported only for quality control/descriptive inspection and is not used for inferential comparisons or hypothesis testing; cross-condition inference in this work is based on embryo-level tempo slopes *m*_anchor_ with embryo-bootstrap confidence intervals (Section 3.11).

Primary tempo readout anchored slope *m*_anchor_ ;We quantify embryo-level developmental tempo using an anchored least-squares slope through the fixed anchor point (*T*0, *T*0). For each start *s*, define the nominal time coordinate

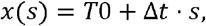

and let

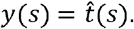

We fit the anchored model

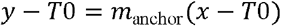

by least squares across starts *s* within the embryo, yielding the closed-form estimate

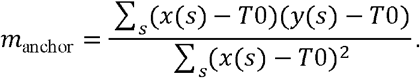

Residuals are computed as

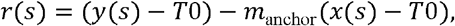

from which we report embryo-level stability metrics: residual RMSE (rmse_resid, hours) and maximum absolute residual (max_abs_resid, hours). Under this definition, *m*_anchor_ <1 indicates a slower tempo relative to the nominal axis, and *m*_anchor_ >1 indicates faster tempo.

Embryo-level quantities (*m*_anchor_, rmse_resid, max_abs_resid) are computed per embryo and used in downstream cross-condition analyses with embryos as the independent statistical units to avoid pseudo-replication [7] (Section 3.11).

### 3.10. SmoothGrad saliency visualization (qualitative)

We use SmoothGrad saliency maps for qualitative inspection of which image regions contribute to clip-level time predictions [23]. For a clip normalized to [0, 1], saliency is defined as the absolute input gradient

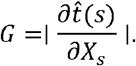

SmoothGrad averages saliency over noisy perturbations 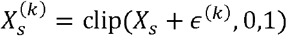 with *ϵ* ^(*k*)^ ∼ *N* (0,*σ*^2^), yielding 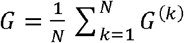. Unless otherwise noted, we use *N*= 20 and *σ* = 0.01.

Per-frame temporal importance is summarized as the spatial mean of for each frame and min–max normalized within each clip for visualization. Heatmap rendering uses percentile stretching, blur, and alpha masking for readability; these steps affect visualization only. Saliency results are reported as qualitative diagnostics and are not interpreted as causal explanations.

### 3.11. Quantification and statistical analysis

Sliding-window clips within an embryo are strongly correlated. Therefore, in all inferential analyses, embryos (not clips/windows) are treated as the independent statistical units to avoid pseudo-replication [7]. Window-level metrics computed over sliding windows are reported for descriptive comparison only and are not used for statistical inference.

Effect size (tempo): Temperature effects are quantified using embryo-level anchored tempo slopes *m*_anchor_ (Section 3.9). We define the tempo effect size as

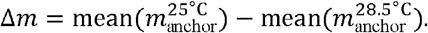

Embryo-bootstrap confidence intervals: Confidence intervals are computed via embryo-level bootstrap resampling [24]. Embryos are resampled with replacement within each condition for *B* = 5000 replicates (seed = 0), and Δ*m* is recomputed for each replicate. A two-sided 95% confidence interval is obtained using the percentile method from the empirical bootstrap distribution of Δ*m*.

Exploratory sample-efficiency (optional): We additionally provide an exploratory power-curve analysis in the released code that retrospectively estimates the probability of detecting Δ*m* ≠ 0 under varying embryo budgets using the same embryo-bootstrap decision rule; this is intended as a planning aid based on the observed dataset distribution rather than a prospective guarantee.

## 4. Results

### 4.1. Overview: clip prediction and embryo-level tempo readouts

Figure 1 summarizes the workflow from clip extraction to embryo-level tempo readouts. The ETF model predicts developmental time from short clips sampled from each brightfield time-lapse sequence. Unless otherwise noted, clips contain *L* = 24 frames sampled at Δ*t* = 0.25 h/frame, and sliding-window inference uses stride *k* = 8, yielding clip-level predictions 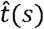 at multiple window starts *s* along each embryo sequence (Section 3.2).

**Fig. 1.**
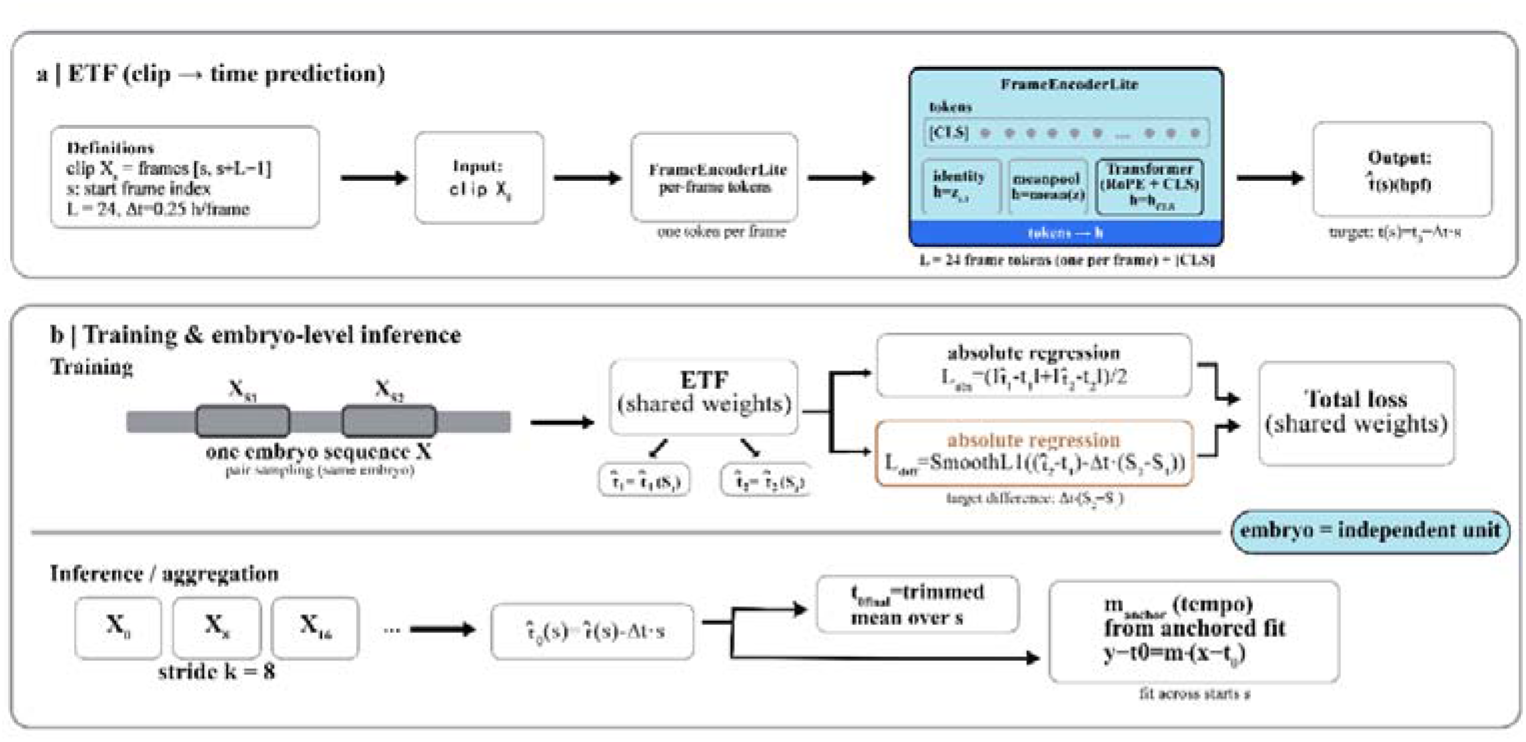
Overview of EmbryoTempoFormer (ETF): clip-to-time prediction, within-embryo temporal-difference consistency training, and embryo-level tempo inference. ETF (the full model) predicts developmental time from clips extracted from zebrafish brightfield time-lapse sequences. Each clip contains L=24 frames sampled at Δt=0.25 h/frame. During training, paired clips from the same embryo are optimized with an absolute regression objective and a temporal-difference consistency regularizer. During inference, sliding-window predictions (stride=8) are converted to - and aggregated robustly to obtain embryo-level readouts, including an anchored tempo slope m_anchor computed from an anchored fit at T0=4.5 hpf. Embryos are treated as independent statistical units for inferential analyses to avoid pseudo-replication [7].

Because sliding-window predictions within an embryo are strongly correlated, downstream analyses summarize clip-level outputs into embryo-level readouts (Section 3.9). We optionally compute an intercept-like diagnostic 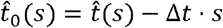 for quality control, but cross-condition comparisons focus on the embryo-level anchored tempo slope *m*_anchor_ and residual-based stability metrics (rmse_resid and max_abs_resid) derived from anchored fits. In all inferential analyses, embryos (not windows) are treated as independent statistical units to avoid pseudo-replication in sliding-window time-lapse evaluation [7] (Section 3.11).

### 4.2. In-distribution ablation at 28.5°C (ID28C5_TEST)

We first evaluate ETF (full model) and three ablations (cnn_single, meanpool, nocons) on the 28.5°C in-distribution test set (ID28C5_TEST; *n* = 22 embryos). Sliding-window inference uses stride = 8, yielding *n* = 484 correlated clip predictions (Section 3.2). The ablations probe progressively stronger temporal modeling assumptions: cnn_single removes temporal context (single-frame baseline), meanpool introduces temporal context but enforces uniform averaging across frames, nocons enables adaptive temporal aggregation via a temporal Transformer but removes the temporal-difference consistency term, and ETF (full) combines Transformer aggregation with within-embryo temporal-difference consistency regularization. This design separates the contributions of temporal context, adaptive temporal aggregation, and consistency regularization.

Under in-distribution conditions, clip-level MAE/RMSE computed over sliding-window clips provide a descriptive summary of time prediction error. Clip-based variants outperform the single-frame baseline, and ETF achieves the lowest clip-level error among the tested models (Fig. 2A; Table 2). Because windows from the same embryo are correlated, these window-level metrics are not used for inferential claims [7].

**Table 2.** In-distribution (28.5°C, ID28C5_TEST) ablation: descriptive clip-level error and anchored-fit consistency metrics (stride = 8). **Table note:** Data are derived from ID28C5_TEST (28.5°C; embryos). Sliding-window inference uses frames, h/frame, and stride, yielding correlated windows. **MAE, RMSE, and R**^**2**^ are computed over pooled window predictions and are reported for **descriptive** comparison only; inferential claims in this work treat embryos as independent statistical units to avoid pseudo-replication [7] (Section 3.11). The tempo slope is computed **per embryo** from an anchored fit at hpf (Section 3.9), and **m_P05 / m_P95** denote the 5th/95th percentiles across embryos. **rmse_resid** and **max_abs_resid** summarize residual scatter of an anchored fit under the model (Section 3.9). All values are rounded to three decimals.

| Model | MAE (h) | RMSE (h) | R2 | m_anchor (mean) | rmse_resid (h) | max_abs_resid (h) | m_P05 | m_P95 |
| --- | --- | --- | --- | --- | --- | --- | --- | --- |
| full (ETF) | 0.746 | 1.029 | 0.993 | 1.009 | 1.006 | 3.777 | 0.963 | 1.045 |
| nocons | 0.811 | 1.119 | 0.992 | 1.014 | 1.066 | 3.802 | 0.962 | 1.048 |
| meanpool | 0.867 | 1.165 | 0.992 | 1.002 | 1.165 | 5.328 | 0.95 | 1.041 |
| cnn_single | 1.063 | 1.513 | 0.986 | 0.984 | 1.462 | 7.116 | 0.933 | 1.022 |

**Fig. 2.**
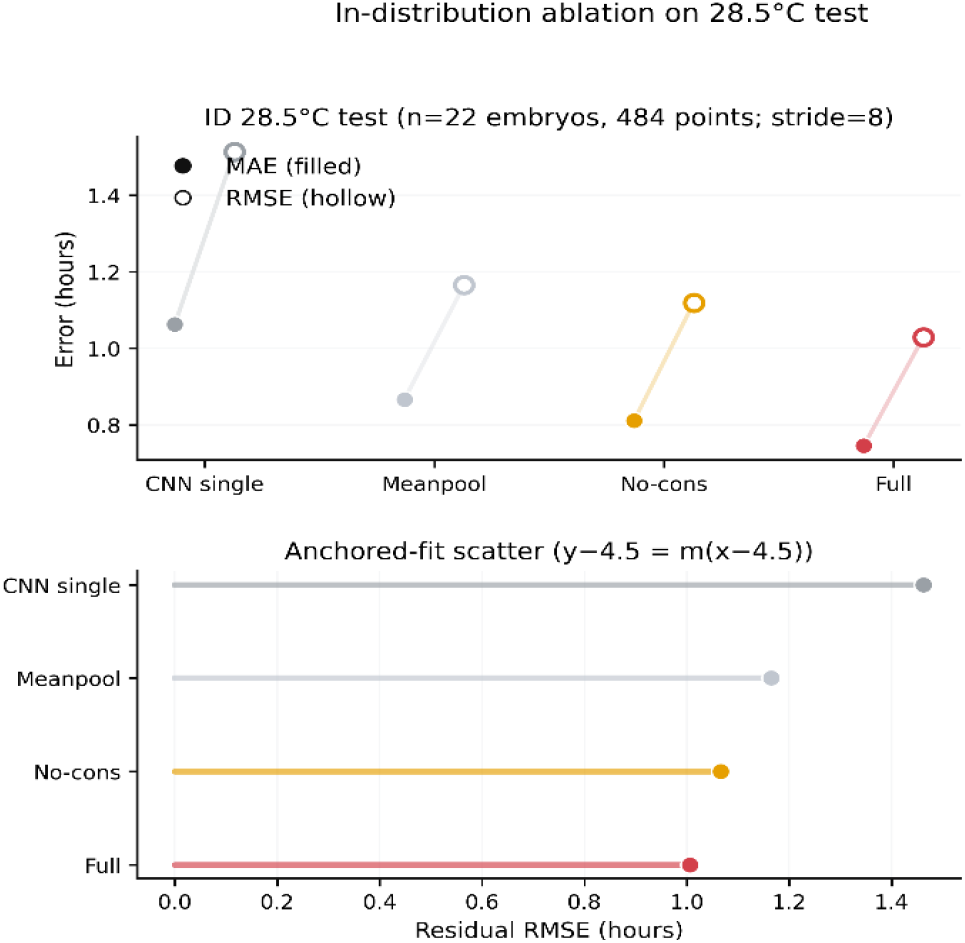
In-distribution ablation on the 28.5°C test set (ID28C5_TEST): descriptive clip-level errors and anchored-fit residual scatter. (A) Clip-level MAE (filled) and RMSE (hollow) computed over all sliding-window clips (stride = 8; correlated windows from embryos). Because windows within an embryo are correlated, these metrics are reported for descriptive comparison only. (B) Anchored-fit residual RMSE (rmse_resid, hours), summarizing how tightly sliding-window predictions align to a single anchored trajectory under the model with hpf (Section 3.9). Models compared: cnn_single, meanpool, nocons, and ETF (full). Numerical values are reported in Table 2.

We therefore emphasize trajectory self-consistency as a key property of time-lapse predictions. Using the anchored-fit framework defined in Section 3.9, ETF yields lower residual scatter (rmse_resid) and improved worst-case residual behavior (max_abs_resid) compared with simpler baselines (Fig. 2B; Table 2). Notably, comparing **nocons** and **ETF** isolates the impact of the temporal-difference consistency regularizer: enabling consistency improves anchored-fit residual behavior while keeping embryos as the inferential units.

### 4.3. External 25°C: tempo distribution and stability (EXT25C_TEST)

We next analyze the external 25°C test set (EXT25C_TEST; *n* = 95 embryos; stride = 8; *n* = 2090 correlated windows). Under temperature shift, the nominal mapping from frame index to hpf is not a ground-truth developmental clock for 25°C; therefore, point-level deviation from the nominal “*y* = *x*” mapping is not interpreted as external-domain accuracy. Instead, we focus on embryo-level tempo and residual-based stability readouts derived from anchored fits (Section 3.9).

Embryo-level tempo slopes at 25°C show tight distributions across embryos. The model-specific 5th–95th percentile ranges of *m*_anchor_ are: ETF 0.664–0.752, nocons 0.673–0.771, meanpool 0.703–0.786, and cnn_single 0.716–0.801 (Fig. 3A; Table 3). These embryo-level slopes provide an interpretable summary of developmental slowdown under temperature shift.

**Table 3.**
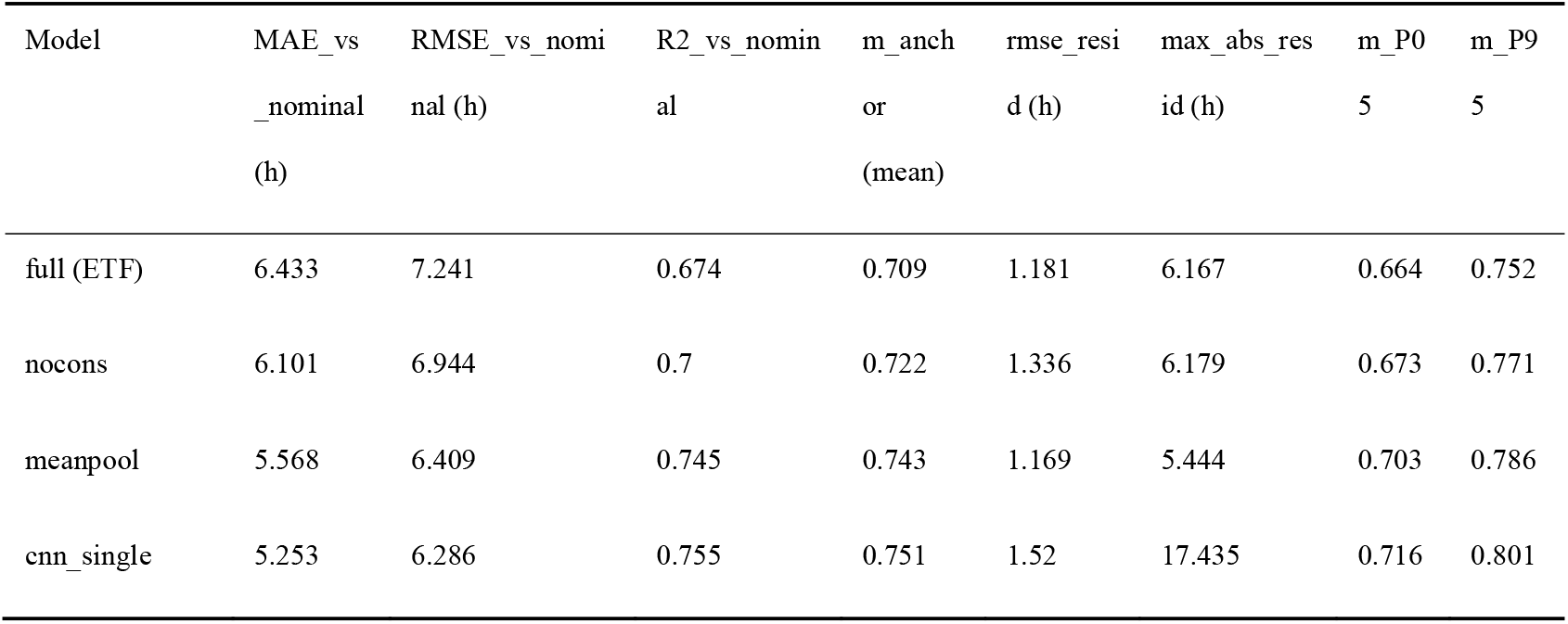
External 25°C test (EXT25C_TEST): descriptive deviation from the nominal time axis and embryo-level tempo/stability summaries (stride = 8). **Table note:** Data are derived from EXT25C_TEST (25°C; *n* = 95 embryos). Sliding-window inference uses **clip length** *L* = 24 frames, **sampling interval** Δ*t* = 0.25 h/frame, and **stride** *k* = 8, yielding *n* = 2090 **correlated** windows. The anchor time is *T*0 = 4.5 hpf. **Point-level (window-level) deviation vs nominal axis (descriptive only):** 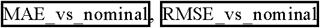, and 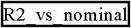 quantify deviation from the nominal mapping *t*(*s*) = *T*0 + Δ*t s* (i.e., deviation from the *y* =*x* line when plotting 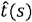 vs. *t*(*s*)). Under a temperature-induced tempo shift, these metrics are reported for descriptive comparison only and are **not** interpreted as external-domain accuracy. **Embryo-level tempo and stability (primary summaries):** For each embryo, tempo is summarized by the anchored slope *m*_anchor_ computed from an anchored fit at *T*0 = 4.5 hpf under (*y*-*T*0) = *m* (*x* - *T*0) (Section 3.9). The table reports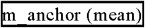 as the **mean across embryos**, and 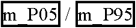 as the 5th/95th percentiles across embryos. Residual-based stability metrics are computed per embryo from the same anchored fit: 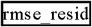 (hours; **mean across embryos**) and outliers. 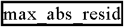 (hours; **maximum across embryos**, i.e., worst-case embryo) to characterize long-tail outliers. Sliding-window predictions are correlated within embryos; inferential analyses treat embryos as independent statistical units to avoid pseudo-replication. Temperature-effect uncertainty is quantified with embryo-bootstrap confidence intervals (Section 3.11) [7]. Values are rounded to three decimals. *(Consistency note for readers: Fig. 3 displays medians and IQRs for visualization; this table reports means (and selected percentiles) for numeric summaries*.)

**Fig. 3.**
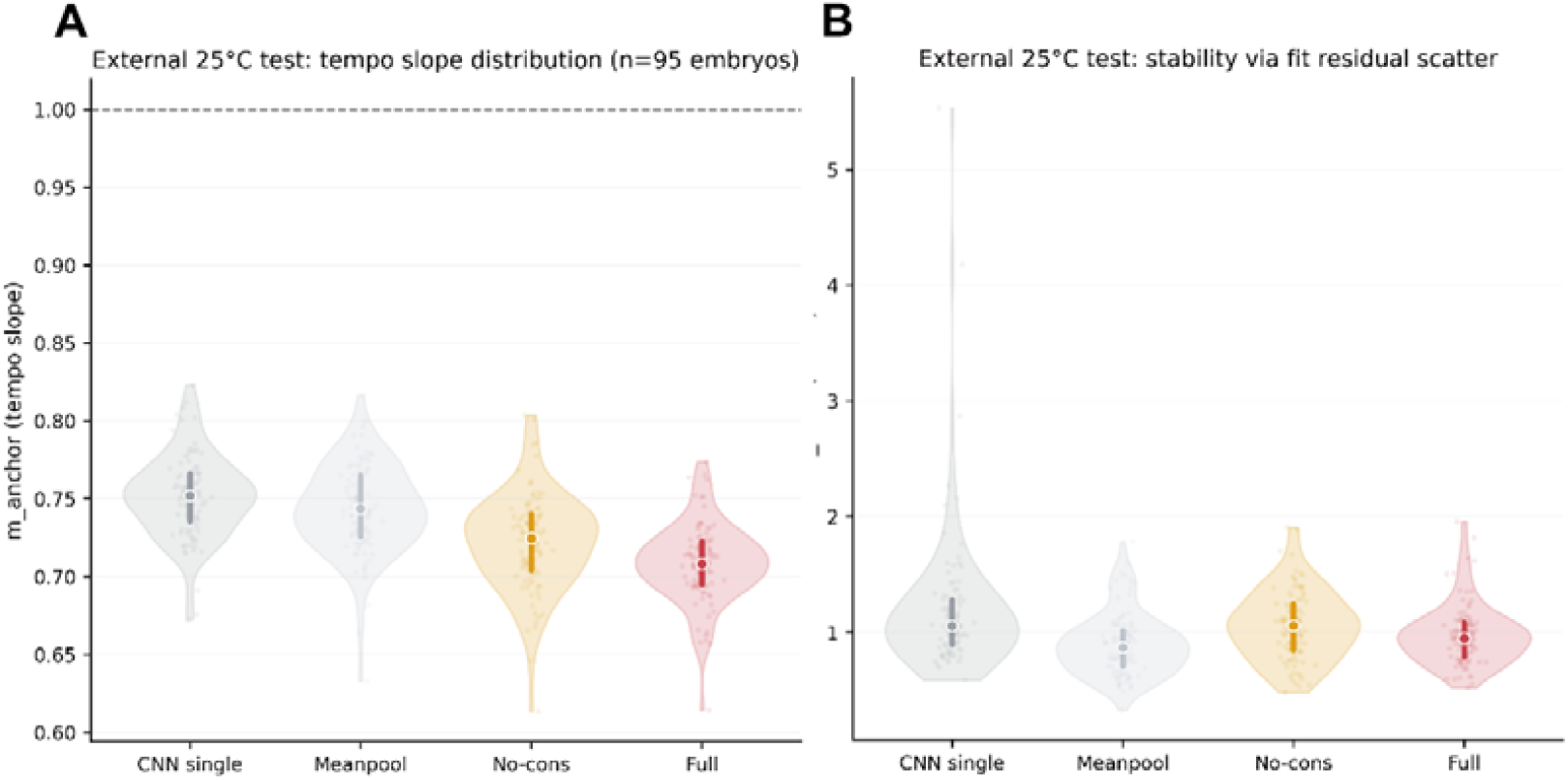
External 25°C test (EXT25C_TEST): embryo-level tempo distribution and residual-based stability across models. (A) Distribution of embryo-level anchored tempo slopes *m*_anchor_ on EXT25C_TEST (25°C; *n* = 95 embryos). Each point represents one embryo. Violins summarize the distribution across embryos; the **white dot indicates the median** and the **thick vertical bar indicates the interquartile range (IQR, 25th–75th percentiles)**. The dashed line at *m* = 1 indicates nominal unit tempo. (B) Distribution of per-embryo anchored-fit residual RMSE (rmse_resid, hours) on the same set, summarizing within-embryo trajectory self-consistency across sliding-window predictions (lower is better). Plot elements are as in (A): points are embryos; violins show density; **white dot = median; thick bar = IQR**. Tempo and residual metrics are computed per embryo from anchored fits at *T*0 = 4.5 hpf under the model (*y*-*T*0) = *m* (*x* - *T*0) (Section 3.9). Sliding-window predictions are correlated within embryos; inferential statements treat embryos as independent statistical units [7]. Numerical summaries are reported in Table 3. Under domain shift, meanpool shows a slightly lower **median** rmse_resid than the full model, consistent with the robustness of uniform temporal averaging to temperature-induced distribution shift.

We further assess trajectory stability using anchored-fit residual metrics (rmse_resid and max_abs_resid) (Fig. 3B; Table 3). These metrics complement tempo slopes by quantifying how coherently sliding-window predictions form a single anchored trajectory within each embryo and how sensitive each model is to outlier windows under domain shift. Notably, meanpool shows slightly lower rmse_resid than the full model (1.169 vs 1.181 h), consistent with the robustness of uniform temporal averaging under distribution shift, whereas cnn_single exhibits substantially heavier long-tail behavior (max_abs_resid = 17.435 h), indicating occasional highly inconsistent windows.

#### Interpretation of Table 3

Under temperature shift, MAE_vs_nominal/RMSE_vs_nominal/R2_vs_nominal quantify deviation from the 28.5°C nominal time axis and are therefore descriptive rather than external-domain accuracy. Consistent with this interpretation, models that infer a stronger slowdown (lower *m*_anchor_) tend to show larger deviation from the nominal axis (e.g., ETF has *m*_anchor_ = 0.709 with MAE_vs_nominal = 6.433 h, whereas cnn_single has *m*_anchor_ = 0.751 with MAE_vs_nominal = 5.253 h).

### 4.4. Temperature effect size and embryo-bootstrap uncertainty

To quantify the temperature-induced tempo shift with statistically valid uncertainty, we compute embryo-bootstrap confidence intervals treating embryos as independent statistical units (Section 3.11). We define the temperature effect size as

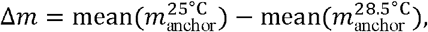

where *m*_anchor_ is the embryo-level anchored tempo slope computed at *T*0 = 4.5 hpf (Section 3.9). Under this definition, negative Δ*m* indicates slower developmental tempo at 25°C.

Across all evaluated models, Δ*m* is negative and the embryo-bootstrap 95% confidence intervals lie entirely below zero (B = 5000; seed = 0), consistent with a robust slowdown under the anchored tempo definition (Fig. 4; Table 4). ETF shows the largest effect magnitude (Δ*m* = -0.300, CI95 [-0.312, -0.288]; Table 4).

**Table 4.** Temperature effect size with embryo-bootstrap 95% confidence intervals (B=5000; seed=0). **Table note:** Effect size definition:, where is the embryo-level tempo slope from an anchored fit at hpf (Section 3.9). Confidence intervals are embryo-bootstrap 95% CIs computed by resampling embryos with replacement within each condition (B = 5000; seed = 0; percentile method; Section 3.11). Embryos are treated as independent statistical units to avoid pseudo-replication from correlated sliding windows [7]. Values are rounded to three decimals.

| Model | Δm | CI95_low | CI95_high |
| --- | --- | --- | --- |
| cnn_single | -0.233 | -0.245 | -0.221 |
| meanpool | -0.259 | -0.273 | -0.244 |
| nocons | -0.292 | -0.304 | -0.279 |
| full (ETF) | -0.3 | -0.312 | -0.288 |

**Fig. 4.**
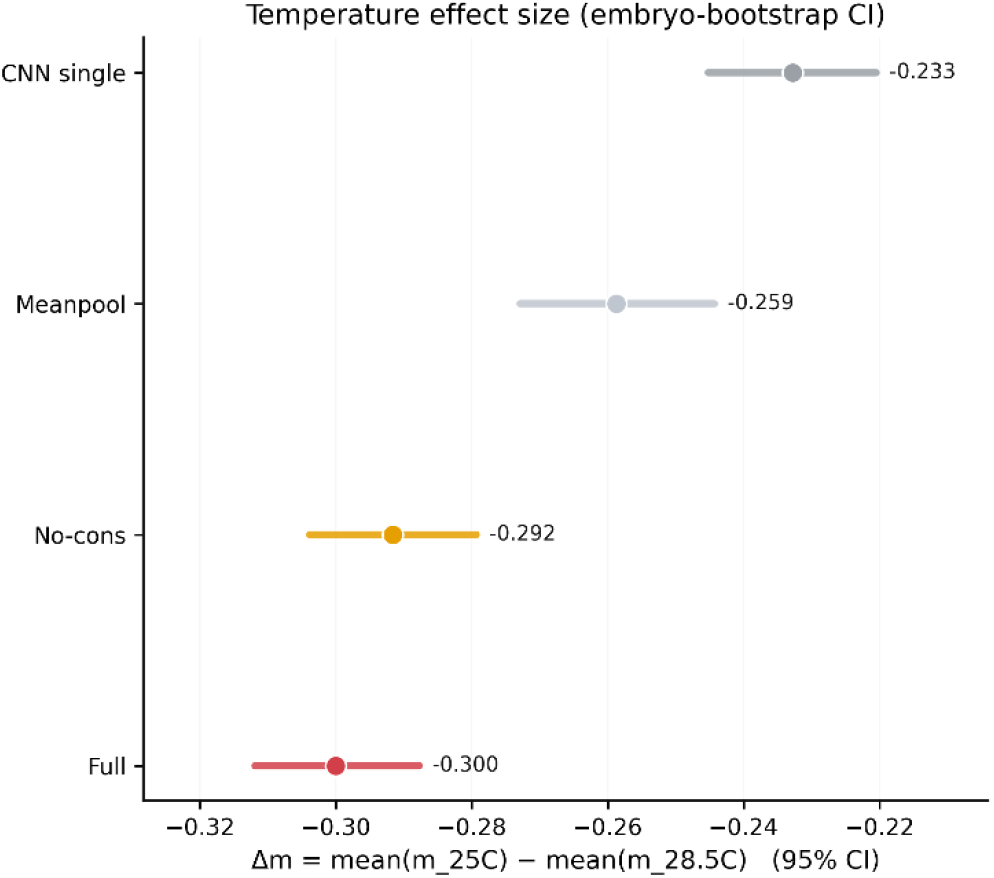
Temperature effect size with embryo-bootstrap 95% confidence intervals (25°C vs 28.5°C). For each model, embryo-level tempo slopes are computed per embryo from anchored fits at hpf under (Section 3.9). The temperature effect size is defined as . Points indicate observed ; horizontal bars show embryo-bootstrap 95% confidence intervals (B = 5000; seed = 0) computed by resampling embryos with replacement within each condition (Section 3.11). Sliding-window predictions are correlated within embryos; embryos are treated as independent statistical units to avoid pseudo-replication [7]. Negative indicates slower developmental tempo at 25°C. Numerical values are reported in Table 4.

We additionally performed a retrospective embryo-level subsampling simulation using the same CI-excludes-zero decision rule; in this dataset, the simulated detection rate saturated at 1.0 by embryos per condition across model variants, and we therefore omit the curve.

### 4.5. Temporal evidence within clips (interpretability)

To provide a qualitative interpretability check and support the intuition that temporal aggregation is not equivalent to uniform averaging, we visualize within-clip temporal importance using SmoothGrad (Fig. 5; Section 3.10). We compute pixel-level saliency as the absolute input gradient 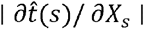 and average it over noisy input perturbations (SmoothGrad). For each frame, we summarize saliency by the spatial mean of the absolute gradient (mean abs-grad) and apply within-clip min–max normalization for visualization, yielding a temporal importance profile (Fig. 5A). Under this normalization, the curve reflects relative within-clip sensitivity across frames rather than absolute saliency magnitude.

**Fig. 5.**
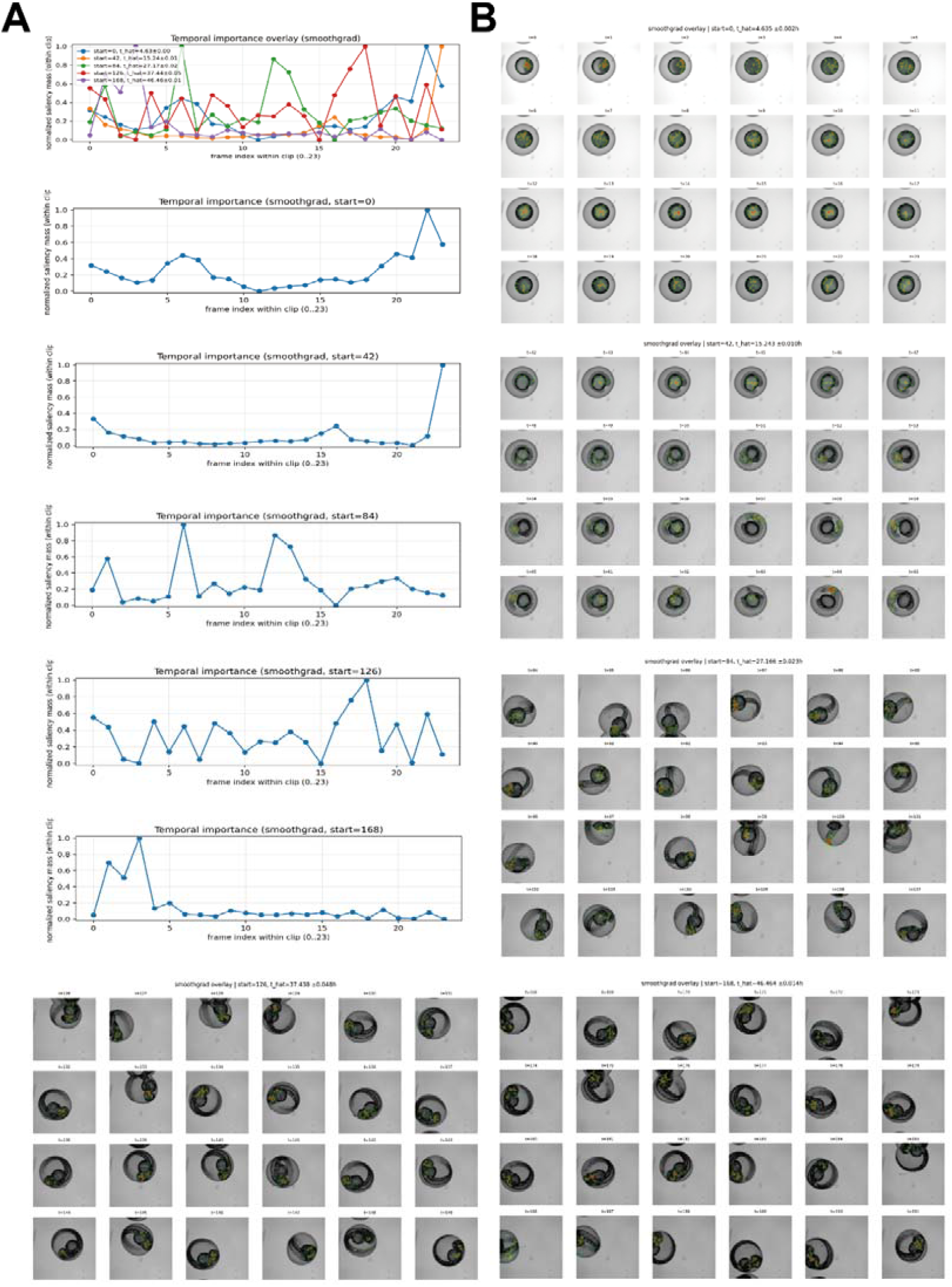
SmoothGrad interpretability suggests non-uniform, phase-dependent evidence usage within 24-frame clips. (A) Within-clip temporal importance profiles for five clip starts (start = 0/42/84/126/168). For each frame, importance is computed as the spatial mean of SmoothGrad saliency 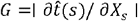, where SmoothGrad averages saliency over *N* = 20 noisy perturbations with *σ* = 0.01 on inputs normalized to [0, 1] . Curves are min–max normalized within each clip for visualization, reflecting relative within-clip sensitivity. (B–F) SmoothGrad heatmap overlays for the corresponding 24-frame clips at the five starts (one panel per start; 24 frames per panel). Heatmaps are visualized with percentile stretching, blur, and alpha masking for readability; these rendering steps affect visualization only. Saliency reflects local sensitivity rather than causal attribution [23].

Across five windows spanning early to late phases (start = 0/42/84/126/168), temporal importance exhibits pronounced non-uniform peaks, and peak locations vary with developmental phase (Fig. 5A). Spatial overlays (Fig. 5B–F) show a phase-dependent shift in sensitivity patterns: early windows (e.g., start = 0; nominal time ≈ 4.5 hpf) emphasize yolk-related morphology and the embryo–yolk boundary; early–mid windows (start = 42 and 84; nominal times ≈ 15.0 and ≈ 25.5 hpf) increasingly highlight axial/caudal (tail) structures alongside persistent yolk-related cues; and mid–late to late windows (start = 126 and 168; nominal times ≈ 36.0 and ≈ 46.5 hpf) show more localized sensitivity around head/eye regions together with yolk-related signal. This qualitative progression is consistent with the increasing availability of localized morphological landmarks over development.

These observations provide an intuitive rationale for moving beyond mean pooling: peaked temporal importance profiles indicate that within-clip evidence is not uniformly distributed across frames, suggesting that a subset of frames may carry higher-information cues for time estimation. Mean pooling enforces equal weighting across frames and may dilute such key-frame signals, whereas Transformer-based aggregation can represent non-uniform evidence usage. We emphasize that SmoothGrad/saliency reflects local sensitivity rather than causal attribution; we therefore use it as a qualitative interpretability check rather than a mechanistic explanation [23].

## 5. Discussion

A central challenge in zebrafish brightfield time-lapse analysis is that sliding-window inference produces many strongly correlated predictions from the same embryo. Treating these window-level outputs as independent samples inflates the effective sample size and constitutes pseudo-replication, leading to overconfident conclusions [7]. In this work, EmbryoTempoFormer (ETF) produces clip-level developmental time predictions 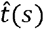 for window start index *s*, while our downstream inference and statistical workflow aggregates temporally correlated window predictions into embryo-level tempo readouts and quantifies uncertainty using embryos—rather than windows—as the independent statistical units. To avoid mixing statistical levels, we explicitly separate (i) window-level error metrics used for descriptive model comparison from (ii) embryo-level tempo and stability readouts used for interpretation and inferential comparisons.

The ablation results disentangle three ingredients for time-lapse staging—temporal context, adaptive temporal evidence fusion, and within-embryo temporal consistency—using a staged comparison (cnn_single → meanpool → nocons → full/ETF ; Table 2). First, adding short temporal context (cnn_single → meanpool) improves descriptive clip-level accuracy on ID28C5_TEST (MAE 1.063 → 0.867 h; RMSE 1.513 → 1.165 h; Table 2), supporting the premise that short temporal windows contain information beyond single-frame morphology. Second, beyond average error, anchored-fit residual behavior summarizes whether overlapping windows align to a coherent within-embryo trajectory—an important property when dense, overlapping predictions are interpreted as a single embryo’s developmental course under correlated sampling [7]. Relative to meanpool, adaptive temporal aggregation (meanpool → nocons) improves both error and residual behavior (MAE 0.867 → 0.811 h; rmse_resid 1.165 → 1.066 h; max_abs_resid 5.328 → 3.802 h; Table 2), consistent with non-uniform evidence fusion reducing inconsistent (outlier) windows. Third, comparing nocons and full/ETF isolates the contribution of the temporal-difference consistency regularizer: enabling consistency improves both average error and anchored-fit residual scatter (MAE 0.811 → 0.746 h; rmse_resid 1.066 → 1.006 h; Table 2), while leaving worst-case residuals largely unchanged (max_abs_resid 3.802 → 3.777 h). Together, these results motivate reporting not only pointwise error but also trajectory-level self-consistency when interpreting dense time-lapse predictions produced by overlapping sliding windows.

Embryo-level tempo readouts become more informative under domain shift, where a nominal clock is not an appropriate external accuracy target. Under temperature change, the nominal mapping from frame index to hpf no longer represents a ground-truth developmental clock for 25°C. Let the nominal coordinate be:

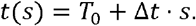

For a given embryo, define *x* (*s*) = *t* (*s*) and 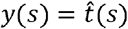. We summarize tempo by the anchored slope *m*_anchor_ from the anchored fit:

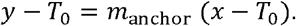

Residuals are defined as:

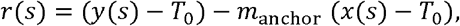

from which we compute embryo-level stability metrics such as residual RMSE (rmse_resid) and the worst-case absolute residual (max_abs_resid). Under this framing, deviation from the nominal mapping (the *y* = *x* line when plotting 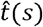 against nominal *t* (*s*)) reflects accumulated deviation from the nominal axis rather than external-domain staging accuracy. This perspective is consistent with recent work emphasizing developmental time and tempo as objects of inference under condition shifts [6]. We therefore focus on embryo-level anchored tempo slopes (reported as m_anchor in tables) and residual-based stability metrics as primary external readouts (Fig. 3; Table 3). On EXT25C_TEST, *m*_anchor_ distributions are tight across embryos (ETF P05–P95: 0.664–0.752; Table 3), enabling interpretable embryo-resolved summaries of developmental slowdown under temperature shift.

Crucially, uncertainty for the temperature effect can be stated at the correct inferential level by treating embryos as independent units. Across all evaluated models, embryo-bootstrap 95% confidence intervals for the effect size

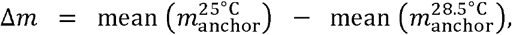

lie entirely below zero (Table 4), consistent with a robust slowdown under the anchored tempo definition. ETF yields the largest effect magnitude (Δ*m* = [-0.300, CI -0.312, -0.2888]; Table 4). Embryo-level bootstrap resampling provides a principled uncertainty estimate when embryos are the independent units and avoids pseudo-replication that would arise from resampling correlated windows [7, 24]. At the same time, external “vs_nominal” metrics should not be interpreted as accuracy: models that infer a stronger slowdown (lower *m*_anchor_) naturally tend to show larger deviation from the 28. 5◦C nominal axis (e.g., ETF: *m*_anchor_ = 0.709, MAE_vs_nominal = 6.433 h; cnn_single : *m*_anchor_ = 0.751, MAE_vs_nominal = 5.253 h; Table 3). Residual-based stability provides complementary information about robustness under shift: meanpool shows slightly lower rmse_resid than ETF at 25◦C (1.169 vs 1.181 h; Table 3), suggesting a mild robustness advantage of uniform averaging under distribution shift, whereas the much larger max_abs_resid for cnn_single (17.435 h; Table 3) indicates occasional highly inconsistent windows when temporal modeling is insufficient. For presentation, Fig. 3 visualizes embryo-level distributions using medians and IQRs, whereas Table 3 reports numeric summaries computed from embryo-level aggregation; these views are complementary.

Together, these results motivate reporting tempo and stability at the embryo level rather than relying on window-level deviation from a nominal clock under condition shifts. Qualitative interpretability further contextualizes why uniform temporal averaging can be limiting. SmoothGrad analyses show non-uniform, phase-dependent temporal importance within 24-frame clips (Fig. 5A), with spatial sensitivity patterns shifting across development (Fig. 5B–F). While saliency reflects local sensitivity rather than causal attribution, these structured patterns are compatible with the intuition that informative morphological landmarks emerge and change over time, supporting aggregation mechanisms that can represent non-uniform evidence usage [23].

Overall, this work complements prior single-image staging approaches such as KimmelNet [5] by leveraging short temporal context and, critically, by centering embryo-resolved inference and embryo-level statistics to avoid pseudo-replication [7]. The approach integrates (i) clip-based modeling to provide local temporal context, (ii) adaptive temporal evidence fusion, (iii) within-embryo temporal-difference consistency to promote coherent trajectories, and (iv) embryo-level tempo readouts with embryo-bootstrap uncertainty to support rigorous, interpretable comparisons across conditions. Limitations are modest but important. External validation here focuses on a single domain shift (temperature), and broader generalization across imaging setups and laboratory protocols remains to be tested. In addition, the anchored tempo definition depends on the chosen anchor *T*_0_; multi-anchor or nonlinear tempo formulations could be explored to assess sensitivity while retaining embryo-level inference. Finally, SmoothGrad is qualitative and method-dependent; it is included as a compatibility check for non-uniform temporal evidence usage rather than a mechanistic explanation [23].

## 6. Conclusions

EmbryoTempoFormer (ETF) is a clip-based CNN–Transformer model for predicting developmental time from zebrafish brightfield time-lapse microscopy. In this study, we pair the ETF model with an embryo-resolved inference and statistical workflow that aggregates temporally correlated sliding-window predictions into interpretable embryo-level tempo and stability readouts, and quantifies cross-condition effects using embryo-bootstrap confidence intervals with embryos treated as independent units to avoid pseudo-replication. By combining lightweight frame encoding, temporal aggregation, and a within-embryo temporal-difference consistency regularizer, ETF improves trajectory self-consistency under dense sliding-window sampling. Together, the ETF model and the accompanying reproducible pipeline (code, scripts, and a Zenodo bundle of processed data, splits, and checkpoints) provide a robust and reproducible basis for high-throughput phenotyping of condition-induced changes in developmental dynamics.

## CRediT authorship contribution statement

Lijiayu Deng: Conceptualization, Investigation, Experimental design, Data analysis, Writing – original draft.

Peiran Lin: Conceptualization, Writing – review & editing.

Luotong Xie: Investigation, Experimental execution.

## Lead Contact

Further information and requests should be directed to and will be fulfilled by the lead contact: Lijiayu Deng.

## Experimental model and subject details

We analyze publicly available zebrafish brightfield time-lapse microscopy data from the BioImage Archive (accession S-BIAD531). Two temperature conditions are considered: 28.5°C (in-distribution) and 25°C (external). Embryo-level dataset splits and exclusions are summarized in Table 1. No new animal experiments were performed in this work.

## Data and Code Availability

Code repository: https://github.com/LijiayuDeng/s-biad531-embryo-tempoformer Zenodo reproducibility bundle (processed arrays, model checkpoints, dataset splits, and checksums): https://doi.org/10.5281/zenodo.18318139

Raw data: BioImage Archive accession S-BIAD531: https://www.ebi.ac.uk/bioimage-archive/galleries/S-BIAD531.html

All quantitative results and figures can be reproduced using the provided scripts (see Reproducibility).

## Declaration of Generative AI and AI-assisted technologies in the writing process

During the preparation of this work, the authors used the OpenAI model gpt-5.2-2025-12-11 to assist with language editing, manuscript organization/formatting, and code suggestions/refactoring for analysis and visualization scripts. After using the tool, the authors reviewed, verified, and edited the material as needed and take full responsibility for the content of the published article.

